# Same-sex sexual behavior and selection for indiscriminate mating

**DOI:** 10.1101/2020.08.12.248096

**Authors:** Brian A Lerch, Maria R Servedio

## Abstract

The widespread presence of same-sex sexual behavior (SSB) has long been thought to pose an evolutionary conundrum^1-3^, as participants in SSB suffer the cost of failing to reproduce after expending the time and energy to find a mate. The potential for SSB to occur as part of an optimal strategy has received almost no attention, although indiscriminate sexual behavior may be the ancestral mode of sexual reproduction^4^. Here, we build a simple model of sexual reproduction and create a theoretical framework for the evolution of indiscriminate sexual behavior. We provide strong support for the hypothesis that SSB is likely maintained by selection for indiscriminate sexual behavior, by showing that indiscriminate mating is the optimal strategy under a wide range of conditions. Further, our model suggests that the conditions that most strongly favor indiscriminate mating were likely present at the origin of sexual behavior. These findings have implications not only for the evolutionary origins of SSB, but also for the evolution of discriminate sexual behavior across the animal kingdom.

---

Empirical observations of same-sex sexual behavior (SSB; i.e., any attempted sexual activity between two or more members of the same sex) in animals are widespread, with evidence of SSB in mammals^5-9^, birds^10-14^, arthropods^15-19^, mollusks^20-22^, echinoderms^23-25^, and other animals^26-30^. Since SSB is traditionally thought to be deleterious, as same-sex matings require energy expenditure but cannot produce offspring, there has been much interest in understanding its origin and maintenance^1-5^. Despite this, there exists no strong theoretical foundation for understanding SSB (but see ^31,32^), resulting in a wide range of untested verbal arguments in the literature^1-5^.

Recently, Monk et al.^4^ challenged the longstanding perspective of SSB as a derived trait, arguing that rather than trying to understand its presence, a more salient question would be to understand its absence. They hypothesize that indiscriminate sexual behavior (i.e., mating without determining the sex of one’s partner) is the ancestral condition, realizing that discriminate sexual behavior (i.e., directing sexual behavior at members of the opposite sex) must evolve through mechanisms controlling sexual signaling and mate choosiness. Of course, the existence of indiscriminate mating as the ancestral condition does not explain its current prevalence^33^. While in some cases (e.g., broadcast spawning and wind pollination) indiscriminate mating predominates as a result of little potential benefit to (or opportunity for) sexual discrimination, it is oftentimes unclear why indiscriminate mating persists.

Building on the perspective of Monk et al.^4^, we argue that selection may actually favor indiscriminate sexual behavior (or prevent the evolution of sexual discrimination) under a wide range of conditions observed in nature. We create a theoretical framework for understanding the conditions that favor indiscriminate sexual behavior and provide a test of whether SSB is likely to result from selection for indiscriminate sexual behavior. We start with a simple optimization model of sexual reproduction, then support this approach with a population genetic model that explicitly tracks evolutionary dynamics. We find that indiscriminate mating is the optimal strategy for many parameter combinations and produce testable predictions about the conditions that favor SSB resulting from indiscriminate mating.

## Optimization Model

We present the optimization model in full in the Supplementary Methods and provide a basic summary of its features here. Our approach explores one of many^34^ potential hypotheses for SSB (that it results from indiscriminate mating) without considering the evolution of same-sex preferences that have evolved in some vertebrates and may result from complex social or genetic interactions (see Table 2 in Bailey and Zuk^1^). As a result, and because our model does not make assumptions consistent with sexual behavior in humans, this study should not be considered in relation to human sexuality. We assume that a population consists of two sexes (the searching sex and the targeted sex), where a proportion σ is of the targeted sex. We make no assumptions about the identity of the sexes and use the terms searching and targeted liberally. For example, if our model were applied to an insect in which males seek females to display to, males would be the searching sex and the females would be the targeted sex.

**Table 1.**
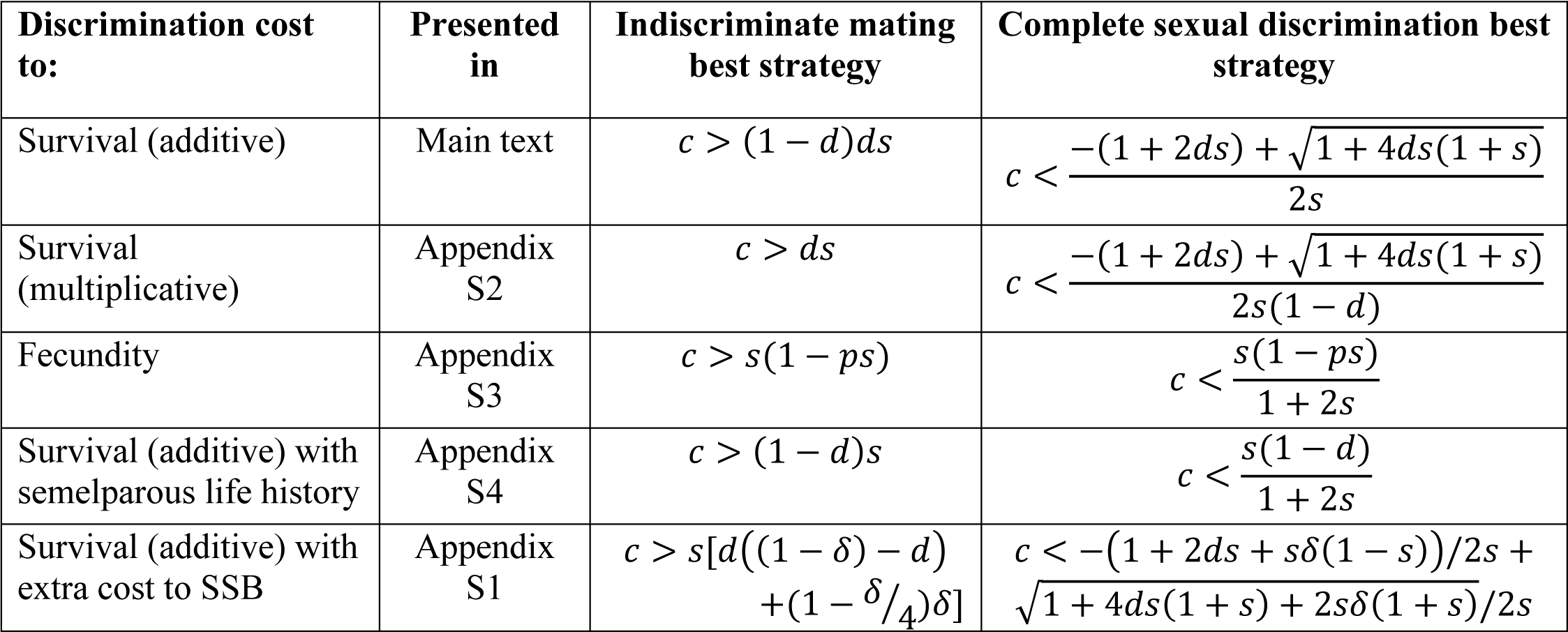
Conditions for no discrimination or complete discrimination to be the optimal evolutionary strategy given an equal sex ratio for each of the models we consider.

| Discrimination cost to: | Presented in | Indiscriminate mating best strategy | Complete sexual discrimination best strategy |
| --- | --- | --- | --- |
| Survival (additive) | Main text | $c > (1 - d)ds$ | $c < \frac{-(1 + 2ds) + \sqrt{1 + 4ds(1 + s)}}{2s}$ |
| Survival (multiplicative) | Appendix S2 | $c > ds$ | $c < \frac{-(1 + 2ds) + \sqrt{1 + 4ds(1 + s)}}{2s(1 - d)}$ |
| Fecundity | Appendix S3 | $c > s(1 - ps)$ | $c < \frac{s(1 - ps)}{1 + 2s}$ |
| Survival (additive) with semelparous life history | Appendix S4 | $c > (1 - d)s$ | $c < \frac{s(1 - d)}{1 + 2s}$ |
| Survival (additive) with extra cost to SSB | Appendix S1 | $c > s[d((1 - \delta) - d) + (1 - \delta/4)\delta]$ | $c < -(1 + 2ds + s\delta(1 - s))/2s + \sqrt{1 + 4ds(1 + s) + 2s\delta(1 + s)}/2s$ |

**Table 2.**
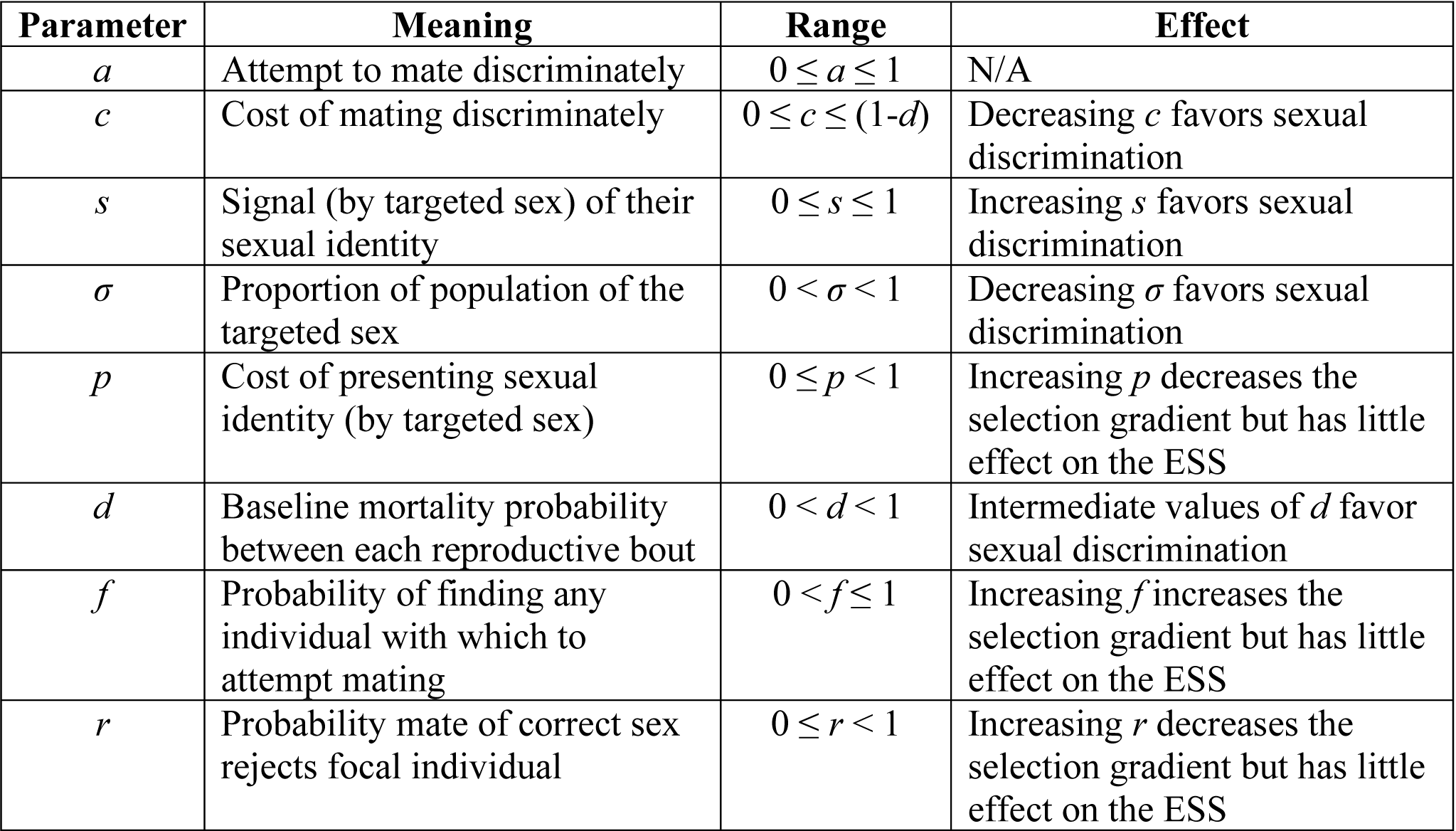
Summary of parameters, their meaning, and their role in the evolution of sexual discrimination from the first model presented with discrimination as an additive cost to survival.

We assume that reproduction occurs in discrete bouts (corresponding to generations) where each member of the searching sex has only one opportunity to mate per bout (an assumption that biases against indiscriminate mating since SSB cannot be corrected for within one reproductive bout). We assume that an individual of the searching sex finds another individual of either sex with which to attempt to mate with probability *f*. The evolutionarily labile parameter of our model *a* controls whether the searching sex attempts to mate discriminately. In particular, *a* is the proportion of bouts in which a member of the searching sex attempts to sexually discriminate. Of course, members of the searching sex can only mate discriminately if they identify some signal (or cue) that an individual is of the opposite sex. We define *s* as the proportion of bouts in which a member of the targeted sex provides a signal of their sexual identity. Then, as shown in the Supplementary Methods, given that a member of the searching sex finds a mate, it will be of the opposite (targeted) sex with probability σ + (1 – σ)*as*. Thus, if members of the targeted sex always signal (*s* = 1) and members of the searching sex always attempt to discriminate (*a* = 1), a member of the searching sex is guaranteed to find a member of the targeted sex. Furthermore, without any signal from the targeted sex (*s* = 0) or any attempt to discriminate from the searching sex (*a* = 0), the probability of finding a mate of the opposite sex is simply the proportion of the population of that sex σ. We discuss the interpretation of *a* and *s* further in the Supplementary Methods.

Even upon finding a mate of the opposite sex the searching sex may be rejected by their potential mate (with probability *r*), in which case they do not reproduce in the reproductive bout. We assume that matings suffer a fecundity cost *p* associated with the sexual signal. Individuals from the searching sex die between reproductive bouts with probability *d* in the absence of sexual discrimination. They also carry an additional survival cost *c* when they attempt sexual discrimination (a search cost), such that a member of the searching sex will survive to the next reproductive bout with probability 1 – (*d* + *ac*).

## Analysis and Results

The model above results in a wide range of parameter space in which indiscriminate mating is an optimal strategy. Specifically, one can derive from this model the expected lifetime reproductive success of a member of the searching sex, *R*_0_. Differentiating *R*_0_ with respect to *a* gives the fitness gradient d*R*_0_/d*a* (see Supplementary Methods). At a given amount of sexual discrimination *a*, the sign of the fitness gradient gives the expected direction of evolution. Values of *a* for which the fitness gradient is 0 are potential evolutionary optima. In analyzing the optimal amount of sexual discrimination, one can determine under what conditions, if any, individuals should attempt to mate indiscriminately. If the optimal strategy is indiscriminate mating, then SSB is expected to be frequent.

Of particular interest is whether indiscriminate mating (*a* = 0) is ever an optimal strategy. We show in the Supplementary Methods that the fitness gradient at *a* = 0 will be negative (and thus sexual discrimination should never evolve) whenever

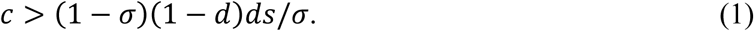

Given a 1:1 sex ratio (σ = 0.5), equation (1) simplifies to *c* > (1 – *d*)*ds*. Equation (1) shows that even under restrictive conditions in which the targeted sex always provides a sexual signal (*s* = 1), the optimal strategy may be to never attempt sexual discrimination. Without sexual signaling (*s* = 0), if there is any cost to attempting to sexually discriminate, sexual discrimination is not expected to evolve. Although this is obvious given the formulation of the model, it formalizes the important point that the origin of sex and the origin of providing signals of one’s sex are not the same. Logically, such cues likely evolved after the origin of sexual reproduction^4^, so our model suggests selection for sexual discrimination was unlikely to follow immediately upon the origin of sex, strengthening the hypothesis that indiscriminate sexual behavior is likely ancestral^4^.

Similarly, the conditions for maximum attempted discrimination *a* = 1 to be the best strategy are derived in the Supplementary Methods and shown in Table 1. If neither condition is met, then an intermediate level of sexual discrimination will evolve (an outcome that occurs in a small but non-trivial portion of the parameter space).

A high cost to sexual discrimination *c* and poor signaling by the targeted sex *s* promotes indiscriminate mating as the optimal strategy (equation (1) and Fig. 1). Sexual discrimination is most likely to evolve when the sex ratio is biased in favor of the searching sex (equation (1)). When the majority of the population is of the targeted sex (1> σ >> 0.5), individuals are more likely to find a member of the opposite sex with which to mate by chance, so attempted sexual discrimination is a worse strategy than when the targeted sex is rare.

**Fig. 1.**
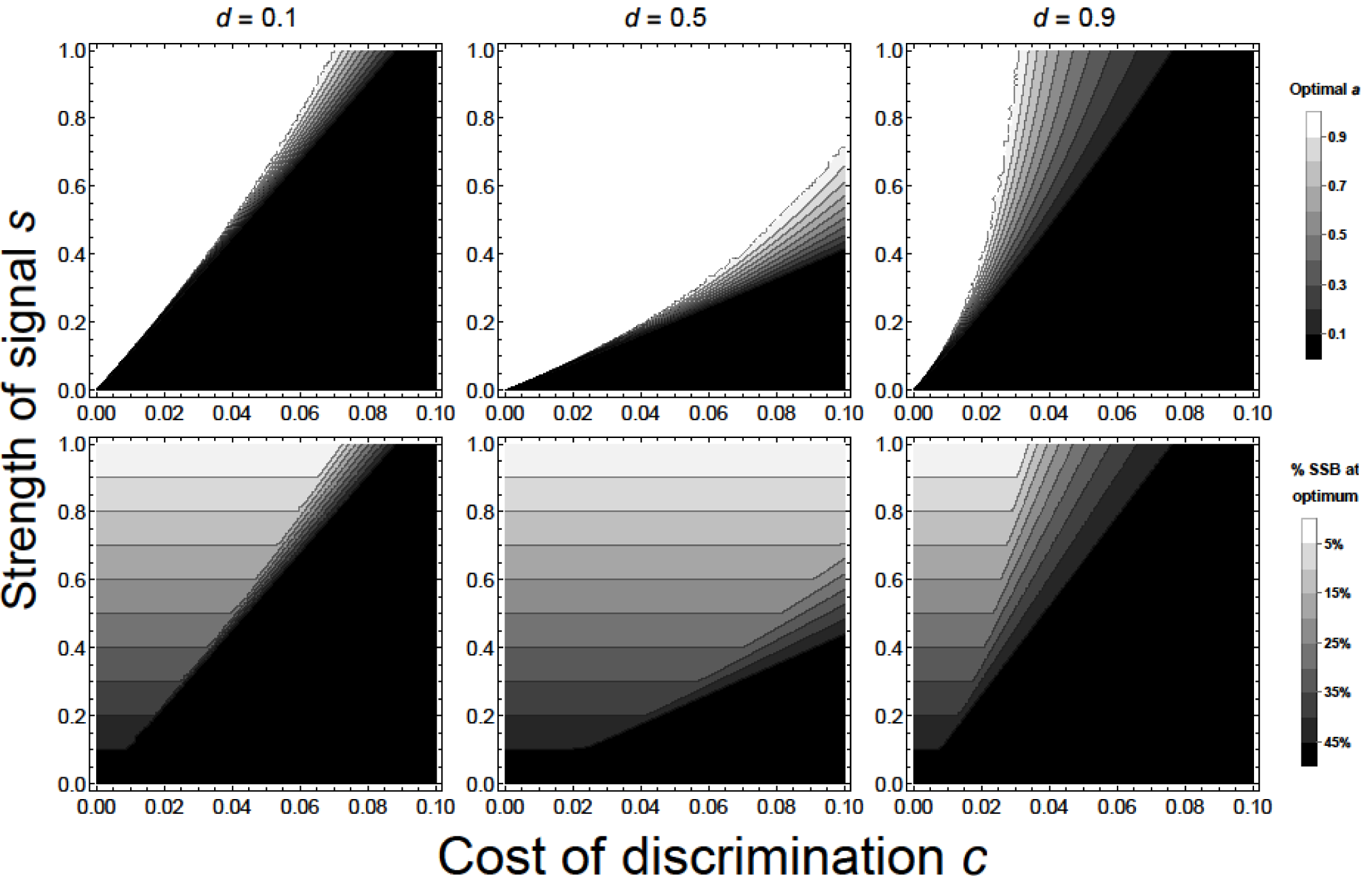
Top row: Optimal discrimination strategies *a* predicted by the optimization model with cost to discrimination *c* on the x-axis, strength of sexual signal *s* on the y-axis, and death rate *d* increasing across columns from left to right. Higher levels of attempted discrimination correspond to lighter shading (white: *a* = 1; black *a* = 0). A wide range of parameter values predict that indiscriminate mating (black) is the best strategy. Indiscriminate mating is favored by increasing the cost of discrimination *c* and decreasing the strength of the sexual signal *s*. Bottom row: proportion of matings expected to be SSB at the evolutionary optimum. Darker values indicate more same-sex matings (black: half of matings are with individuals of the same sex at the optimum; white: no matings are with individuals of the same sex if behaving optimally). Other parameters: proportion of the population of the targeted sex σ = 0.5, probability of finding any individual with which to attempt to mate *f* = 1, probability of being rejected by a potential mate *r* = 0, cost of sexual signal *p* = 0.

Interestingly, an intermediate death rate *d* favors the evolution of sexual discrimination (Fig. 1). When death is rare (small *d*), members of the searching sex are expected to have many reproductive opportunities in their lifetime. Under these conditions, the best strategy is to live as long as possible by not attempting to sexually discriminate. The cost of SSB in this case is low because one failed mating due to SSB will likely be made up for by chance later in life. On the other hand, when *d* is high, members of the searching sex are unlikely to ever mate. In this case, they cannot afford to pay any additional cost and their optimal strategy is to mate indiscriminately and rely on luck. Of course, indiscriminate mating will result in SSB being common (Fig. 1, bottom row).

Although they do not affect the optimal level of discrimination, increasing the cost of sexual signals *p* and the probability of mate rejection *r* and decreasing the probability of finding any individual *f* cause the selection gradient to approach 0 (i.e., weaker selection; Movie S1 and Table 2 show the effect of each parameter). If indiscriminate mating is ancestral, these conditions are more conducive to the transient maintenance of indiscriminate sexual behavior by reducing the efficacy of selection and making the stochastic loss of discriminate mating more likely. Thus, discriminate mating is less likely to evolve in sparse populations (low *f*) or when the targeted sex is choosy or the searching sex competitive (high *r*).

We test the generality of our results by modifying our assumptions to allow same-sex matings to carry an additional cost (Supplementary Appendix S1), to include mortality from different sources acting multiplicatively (Supplementary Appendix S2), to assume the cost to sexual discrimination is due to fecundity as opposed to survival (Supplementary Appendix S3), and to assume a semelparous life history (Supplementary Appendix S4). We show the conditions for no or complete sexual discrimination to evolve given these assumptions in Table 1. We consider the existence of additional costs to SSB as an extension since while such costs have been found^35^ (and are often suggested^3^), other studies fail to support that such costs exist^36,37^. Our primary results are robust to all of these changes, with each version of the model predicting an appreciable region of parameter space for which indiscriminate mating is the optimal strategy. Of course, assuming that SSB carries explicit costs (in addition to the opportunity costs implicit in the above analysis) results in more restrictive conditions for sexual discrimination to evolve, although small costs to SSB have only small impacts on the model outcomes. The only qualitative differences between the model versions occur at high death rates *d* when mortality is multiplicative, high signaling costs *p* when discrimination cost is to fecundity, or low death rates *d* when the searching sex is semelparous. Qualitative outcomes of the models are compared in Table 3.

**Table 3.**
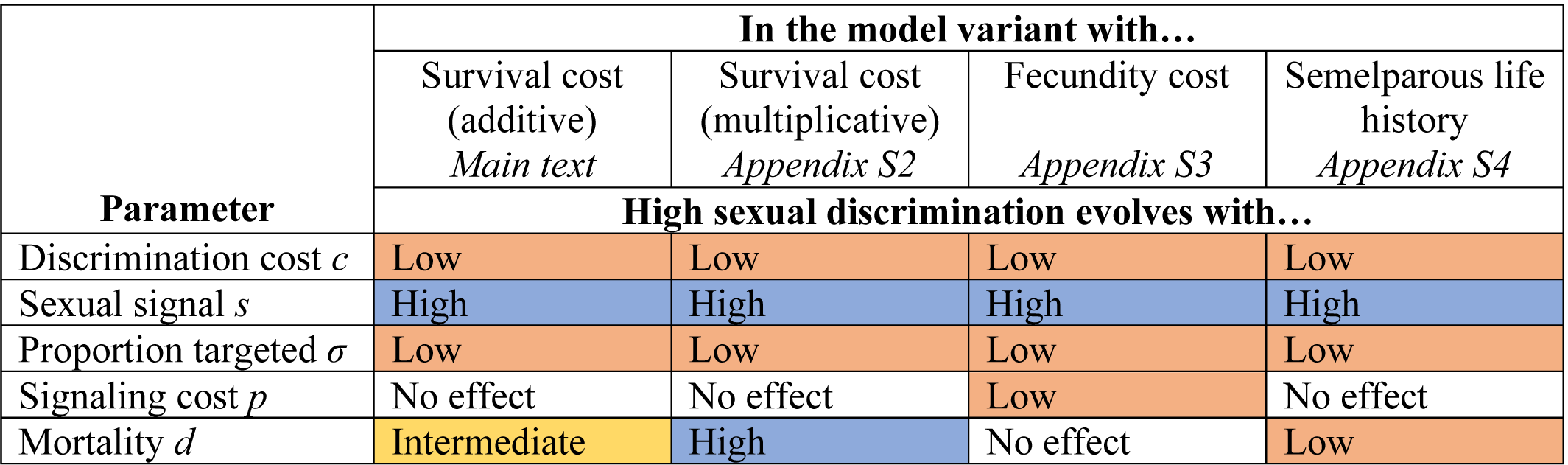
Summary of how key parameters qualitatively influence optimal sexual discrimination in each model variant considered. The four distinct variants are shown as columns and are presented in the location indicated by italics. The information in the table should be read as in the following example. For the top left cell in the main body of the table: in the model variant where there are additive survival costs to discrimination, high sexual discrimination evolves with a low discrimination cost *c*.

| Parameter | In the model variant with... |  |  |  |
| --- | --- | --- | --- | --- |
|  | Survival cost<br>(additive)<br>*Main text* | Survival cost<br>(multiplicative)<br>*Appendix S2* | Fecundity cost<br>*Appendix S3* | Semelparous life<br>history<br>*Appendix S4* |
|  | High sexual discrimination evolves with... |  |  |  |
| Discrimination cost $c$ | Low | Low | Low | Low |
| Sexual signal $s$ | High | High | High | High |
| Proportion targeted $\sigma$ | Low | Low | Low | Low |
| Signaling cost $p$ | No effect | No effect | Low | No effect |
| Mortality $d$ | Intermediate | High | No effect | Low |

## Dynamical Model

Although our optimization model is analytically tractable and clarifies costs and benefits, lifetime reproductive success is not necessarily maximized by selection^38^. As such, we also build a single-locus population genetic model with haploid genetics and overlapping generations that makes similar assumptions to the approach above. Importantly, the population genetic model extends the optimization approach by incorporating frequency dependence and allowing the sex ratio to change naturally from feedbacks with mortality due to discrimination costs. Since *p, r*, and *f* play no role in this framework, they are ignored. We still assume a background mortality of *d* afflicts both sexes and a survival cost of attempted discrimination of *ac* is suffered only by the searching sex. The probability of finding a mate of the opposite sex is still σ + (1 – σ)*as*, but now the sex ratio σ emerges from the model. We use successive invasions to determine the evolutionarily stable values of attempted sexual discrimination *a*. Although not analytically tractable, this model makes no assumptions *a priori* about the quantity that selection maximizes.

The results from this population genetic model align strikingly well with the optimization approach, with the range of conditions under which indiscriminate mating is uninvadible being practically identical between approaches (Fig. S1). A stable polymorphism identified by mutual invasibility only occurs in about 2% of 10,000 randomly generated parameter combinations and is especially common at low or high death rates *d* and strong sexual signals *s*. This model shows that not attempting to discern the sex of potential mates can be a convergent stable evolutionary optimum.

## Implications

SSB is often considered a result of mistaken identity^39-41^, as is suggested to account for about 80% of reported cases in arthropods^3^. Our model provides an evolutionary perspective on this mistaken identity hypothesis, suggesting that poor sex identification could actually occur as an optimal strategy. This evokes hypotheses that SSB may result from a mating strategy of attempting copulation with any encountered conspecific due to low probability of encounter^21^ or low costs to SSB^22,30^. The costs of missing an opportunity to mate and of attempting SSB have been discussed^42-44^ in the context of the acceptance cost threshold hypothesis^45^—a general theory suggesting that erroneous associations (e.g. between mates or cooperative partners) become more likely with poor discrimination ability and low costs to mistaken associations. We provide a formal application of this hypothesis to SSB and show which conditions favor indiscriminate mating.

It was argued by Parker^46^ that six evolutionary transitions (the “sexual cascade”) drove unicellular asexual organisms to become behaviorally-complex, sexual organisms. One such transition is the evolution of the movement of males toward females before sperm release during external fertilization (an example of sexual discrimination referred to as “female targeting”^46^). To our knowledge, Parker’s model^46^ is the only study addressing whether sexual discrimination evolves. Direct comparison between Parker’s model and our model is difficult, but we seem to find more restrictive conditions for the evolution of sexual discrimination, which can be attributed to 1) imperfect signaling (our *s*) of the targeted sex (as is likely at the origin of sexual reproduction^4^) and 2) a search cost^47,48^ (our *c*) for attempting to sexually discriminate (instead of a tradeoff with gonad expenditure^46^). These models are complementary; our model applies to cases not considered by Parker^46^ such as SSB in species with internal fertilization (e.g. insects^3^) or species with limits on their ability to find and identify mates (e.g. those with search costs, poor signals of sexual identity, and deep-sea species^,21,23,24^).

It is interesting to consider how the predictions of the model relate to the conditions expected at the origin of sexual behavior. Echinoderms are likely good proxies for such animals^4,46^, supported by their position as an outgroup to chordates (where most complexity in sexual behaviors arise). Consistent with the model’s predictions for species that mate indiscriminately, long-lived adults are common in echinoderms^49,50^. Additionally, it is reasonable to expect that cues to determine sex in echinoderms are relatively limited both because visual cues cannot be relied upon and there exists little evidence in this taxon for chemical cues for sex-specific recognition from a distance^51,52^. Indeed, multiple studies suggest some echinoderm species form mating pairs without consideration for sex^23,24,53^. This suggests that if indiscriminate sexual behavior is the ancestral condition^4^, sexual discrimination was unlikely to have evolved readily.

This model relates to previous work on mate choice in which there can be a direct cost of mating with one category of individuals versus another^54^, in finding that costs can prevent mating preferences from evolving. In fact, all cases where there are direct viability or fecundity benefits to choosing one type of mate are also somewhat analogous. However, the costs of indiscriminate mating in the current model are much higher than in many other cases with direct benefits, as SSB results in a mating which cannot produce any offspring at all. The mechanisms operating here are most similar to the evolution of preferences for conspecifics, where mating with a heterospecific produces no viable hybrids. In both cases, costs of discrimination will trade off against the peril of producing no offspring. In the current context, the unexpected consequence is that SSB often results.

By showing that there are a broad range of conditions for which indiscriminate mating can be an optimal strategy, we extend recent work^4^ suggesting the evolutionary origins of discriminate sexual behavior as a new and fruitful area of research. Our model provides an important proof-of-concept regarding whether indiscriminate mating can be an optimal evolutionary strategy and what conditions facilitate its evolution. One important result from this modeling exercise is that sexual discrimination can be favored by either low, intermediate, or high mortality rates depending on other features of the system (Table 3). As such, mortality rates alone are unlikely to predict whether indiscriminate mating is an optimal strategy. Costs to discrimination *c* and strengths of sexual signals *s* are more likely candidates for the drivers of indiscriminate mating, but their values in natural populations are unknown. Attempts to measure these (or related) parameters are important gaps to fill in determining whether SSB results from selection for indiscriminate mating in nature. Specifically, our model leads to the predictions that species that mate indiscriminately have high costs to discrimination, search costs to survival rather than fecundity (since this results in more stringent conditions for sexual discrimination, Supplementary Appendix S3), and subtle differences between the sexes. In this way, our model suggests which features of organisms are likely to predispose them to indiscriminate mating, providing a guide future work to determine how widespread selection favoring indiscriminate mating is in nature

## Acknowledgements

We thank Michael Moore and Max Lambert for comments on early versions of the manuscript and Aimee Deconinck for suggesting the name “targeted” sex.

## Author Contributions

BAL conceived of the project and the optimization models. BAL and MRS designed the population genetic models. BAL led the writing on the manuscript with input from MRS.

## Data Availability

All analyses can be reproduced by the files in the supplementary information.

