## Supplemental materials for "Same-sex sexual behavior and selection for indiscriminate mating"

*Supplementary Materials*

Brian A. Lerch and Maria R. Servedio

Contents:

| | |
|---|---|
| Pg. 2 | Supplementary Figure S1 |
| Pg. 3 | Supplementary Methods |
| Pg. 12 | Supplementary Appendix S1: Additional cost to SSB |
| Pg. 16 | Supplementary Appendix S2: Multiplicative mortality |
| Pg. 20 | Supplementary Appendix S3: Fecundity cost to attempted discrimination |
| Pg. 25 | Supplementary Appendix S4: Semelparous life history |

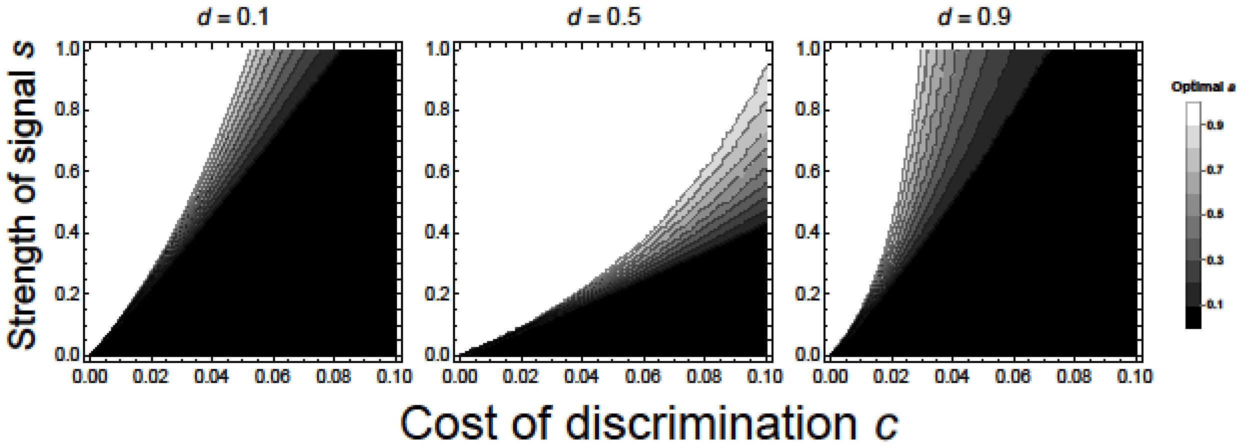


**Fig. S1.** ESS level of attempted sexual discrimination from the population genetic model (lighter colors indicate higher attempted discrimination; white: $a = 1$; black: $a = 0$). That this aligns so well with the top row of Fig. 1 shows that there exist dynamical scenarios where the strategy favored by selection will be to forsake any attempt to determine the sex of potential mates.

**ATTACHED SEPARATELY**

**Movie S1.** Supplementary movie showing the fitness gradient as a function of attempted discrimination $a$ changing with each parameter. The default setting for each parameter is $c = 0.1$, $s = 1$, $\sigma = 0.5$, $d = 0.8$, $f = 1$, $p = 0$, and $r = 0$. Each parameter is allowed to vary between 0 and 1 except for $c$ which is only allowed to vary between 0 and 0.2 and $d$ which is allowed to vary between 0 and 0.9. Evolutionary optima occur wherever the line crosses the x-axis (and has negative slope). If the line is always positive, the optimal strategy is to always discriminate. If the line is always negative, the optimal strategy is to never attempt sexual discrimination.

### Supplementary Methods

#### Optimization model

We assume that a population consists of obligate sexually reproducing organisms which belong to either the searching sex or the targeted sex. As discussed in the main text, our model makes no further assumptions about sexual identity and these terms are meant to be used liberally. We form a simple model for the expected lifetime reproductive success of members of the searching sex (which can sexually discriminate). We do not explicitly consider the lifetime reproductive success of the targeted sex here, as they do not express the ability to sexually discriminate (but they are accounted for in the population genetic model, below).

We assume that reproduction occurs in discrete bouts (corresponding to generations) and that individuals have only one opportunity to reproduce per breeding bout. Considering that the cost of failure to sexually discriminate is that some individuals will not get to reproduce as a result of indiscriminate SSB, this assumption biases against SSB, as it maximizes the cost that individuals pay relative to how often they breed. We assume that the probability that a member of the searching sex (the sex with the opportunity to sexually discriminate) finds an individual of either sex with which to attempt to mate during a breeding bout is $f$. Then, the probability that they find an individual of the opposite sex while randomly searching is the proportion of the population that is of the targeted sex $\sigma$. The strength of attempted sexual discrimination $a$ increases the likelihood that the individual found is of the correct sex. Of course, discrimination can only occur if individuals of the targeted sex provide some signal $s$ of their sexual identity. Such a signal could take on many forms, such as a chemical cue or visual dimorphism. Specifically, we assume that a member of the searching sex attempts to discriminate with probability $a$ and a member of the targeted sex signals with probability $s$. These need not be taken to literally mean that the attempt to search and sexual signal are binary. Rather, $a$ and $s$ are the proportion of time that the attempt to discriminate or sexual signal has an effect. It is worth noting that the parameters $a$ and $s$ can also be interpreted as a strength of attempted discrimination and signal, respectively. With these definitions, given that a member of the searching sex finds a conspecific, the probability that they will be of the opposite (targeted) sex $\zeta$ is

$$\zeta = (1-a)(1-s)\sigma + (1-a)s\sigma + a(1-s)\sigma + as = \sigma + (1-\sigma)as. \tag{1}$$

Here, the first term ($(1-a)(1-s)\sigma$) represents failures to signal and attempt to discriminate simultaneously, the second term ($(1-a)s\sigma$) represents a sufficient signal but no attempt to discriminate, and the third term ($a(1-s)\sigma$) represents an attempt to discriminate but no signal. In each of these cases, the proportion of opposite-sex matings is given by the sex ratio (hence each term is multiplied by $\sigma$). Finally, the fourth term is when both the signal and attempt to discriminate occur, in which case all matings are between individuals of the opposite sex (so this term can be thought of as being multiplied by unity). Complete sexual discrimination occurs if, and only if, $a = s = 1$, and no sexual discrimination occurs if $a = 0$ or $s = 0$. We further assume that even if a member of the searching sex finds a member of the correct sex they are rejected as a mate with probability $r$. Again, "rejection" should be used liberally, as this could not only reflect a conscious choice to reject a mate, but could also correspond to "rejection" by sperm competition.

We assume that no individual variation in reproductive output from a successful mating exists. Of course, if an individual of the searching sex attempts to mate with a member of their own sex, then that mating will produce no offspring. We allow for the possibility that providing sexual signals comes at a relative reproductive cost $p$ for the targeted sex. This cost is considered to be a result of energy expenditure lost by sexually signaling. As such, each mating produces $1 - sp$ offspring. Thus, individuals that sexually signal have a fitness of $1 - p$ relative to those that do not signal. Note that $0 \leq p < 1$.

We assume that individuals of the searching sex have a baseline death probability of $d$ in between each reproductive bout. We further assume that attempting sexual discrimination comes with an additional survival cost $c$ such that individuals that attempt to sexually discriminate suffer an additional probability of death $c$. This cost could correspond, for example, to increased energy expenditure or conspicuousness to predators in attempting identify mates of the opposite sex. Then, the probability of surviving to the next breeding bout $\rho$ is

$$\rho = 1 - (d + ac), \tag{2}$$

where $0 \leq c < 1 - d$ to guarantee the cost is valid for any value of $a$.

From here, it is easy to compute the expected lifetime reproductive success $R_0$ of a member of the searching sex. $R_0$ is a infinite sum of powers of $\rho$ given by

$$R_0 = \sum_{t=1}^{\infty} \zeta(1 - sp)\rho^t. \tag{3}$$

This converges to

$$R_0 = \frac{\rho}{1 - \rho}\zeta(1 - sp). \tag{4}$$

Substituting back the original parameters, we get

$$R_0 = \frac{1 - (d + ac)}{d + ac}(\sigma + (1 - \sigma)as)(1 - r)(1 - sp), \tag{5}$$

which can be thought of as a function of sexual discrimination $a$. We can use basic calculus tools to find the extrema of $R_0$. These are potential evolutionary singular points.

In particular, we can compute the fitness gradient $\frac{dR_0}{da}$ which has a sign that gives the direction of evolution at a given value for $a$. The fitness gradient is

$$\frac{dR_0}{da} = -\frac{f(1-sp)(1-r)[(1-\sigma)s(a^2c^2+2acd+d^2-d)+c\sigma]}{(d+ac)^2}. \tag{6}$$

Solving for where $\frac{dR_0}{da} = 0$ gives potential evolutionary optima. Although tractable, this is not particularly enlightening. Still, equation (6) gives the exact form of the fitness gradient and analyzing it numerically allows for a determination of optimal evolutionary strategies. In particular, the top row of Fig. 1 is made by calculating the values of $a$ for which $\frac{dR_0}{da} = 0$ and ensuring that this is a fitness maximum and not a fitness minimum (i.e., $\frac{d^2R_0}{da^2} < 0$).

It is possible to derive conditions from equation (6) for the evolution of sexual discrimination. The sign of the fitness gradient evaluated at $a = 0$ is of particular interest. Whenever $\frac{dR_0}{da}\big|_{a=0} < 0$, selection acts against attempts to sexually discriminate in the absence of discrimination. In other words, the values for which $\frac{dR_0}{da}\big|_{a=0} < 0$ are vital as these are the conditions for completely indiscriminate mating to be selected for. Fortunately, this can easily be derived. First, notice that the first three terms in the numerator are always positive. Then, because of the negative sign out front, the fitness gradient will be negative whenever the last term in the numerator is positive. That is, whenever

$$(1-\sigma)s(a^2c^2+2acd+d^2-d)+c\sigma > 0. \tag{7}$$

Evaluating equation (7) at $a = 0$ and rearranging gives the condition

$$c > \frac{(1-\sigma)(1-d)ds}{\sigma}. \tag{8}$$

The implications and interpretation of equation (8) are discussed in the main text. Note that, if $a = 0$, then the searching sex suffers no additional cost, so it could be argued that the sex ratio would be expected to be equal under these conditions. Assuming an equal sex ratio, equation (8) simplifies to

$$c > (1-d)ds. \tag{9}$$

Thus, given an equal sex ratio, when equation (9) is satisfied indiscriminate mating is an evolutionary optimum.

Finally, it is also desirable to know what conditions favor maximal attempts to discriminate. In other words, when is the fitness gradient positive at $a = 1$. Again assuming an equal sex ratio (for simplicity), reversing the inequality from equation (7), and evaluating at $a = 1$ gives this condition as

$$c < \frac{-(1 + 2ds) + \sqrt{1 + 4ds(1 + s)}}{2s}. \tag{10}$$

Whenever equation (10) is satisfied, sexual discrimination is expected to evolve to completion. Note that our assumption of an equal sex ratio when $a = 1$ (invoked only to find equation (10)) is not expected to be satisfied due to search costs of sexual discrimination; however, decreasing the proportion of the population of the targeted sex will only make the conditions for complete discrimination to evolve more restrictive. Of course, if neither equation (8) nor equation (10) is satisfied, then there is an evolutionary optimum for an intermediate value of $a$ (Fig. 1, main text).

### Population genetic model

Although the approach described above easily lends itself to analytical techniques, it relies on expected lifetime reproductive success being the target of selection. This is not necessarily true, in general. To get a better sense of the realism of the optimization model, we also developed a population genetic model that makes similar assumptions. Due to the similarity of these models, we note that the definitions of parameters are the same as was given above and avoid redefining them here.

We will consider a scenario where attempted sexual discrimination is controlled by a single locus with two alleles $A_1$ and $A_2$ that attempt to sexually discriminate a proportion of the time $a_1$ and $a_2$, respectively. The proportion of the (haploid) population carrying the allele $A_i$ is $x_i$, where the census is taken at the beginning of each reproductive bout. We will make the standard assumption that the population size is fixed between time steps so that $x_1 + x_2 = 1$, although the $x_i$ may not sum to 1 at other stages of the life cycle and is thus only a true genotype frequency at the census point. In this scenario, it is important to differentiate between the searching sex and the targeted sex, even though the $A$ allele is only expressed by the searching sex. We denote the targeted sex with the superscript $t$ and the searching sex with the superscript $s$. As an example, the proportion of the population that consists of the searching sex that carries the $A_1$ allele is denoted by $x_1^s$.

The first step in the life cycle is death. A proportion $d$ of members of the targeted sex die prior to breeding. The searching sex suffers an additional cost due to attempt to discriminate. In particular, a proportion $d + a_i c$ of $x_i^s$ die prior to breeding. The number of surviving members of each sex will be denoted with a prime $'$.

Following the assumptions made for the optimization model, we next form mating pairs. Because the searching sex is limiting, the number of pairs formed cannot exceed the number of individuals left from the searching sex. Once again $\zeta = \sigma + (1 - \sigma)as$ (derived above) is the probability that a member of the searching sex finds an individual of the opposite sex to mate with. Note that now the sex ratio

$$\sigma = \frac{x_1^{t'} + x_2^{t'}}{x_1^{t'} + x_2^{t'} + x_1^{s'} + x_2^{s'}} \tag{11}$$

emerges naturally from our model. With these assumptions in mind the number of matings between a member of the searching sex with the $A_i$ allele with a member of the targeted sex with the $A_j$ allele $p_{ij}$ is

$$p_{ij} = x_i^{s'}(\sigma + (1-\sigma)a_1 s)\frac{x_j^{t'}}{x_1^{t'} + x_2^{t'}}. \tag{12}$$

The number of individuals of the searching sex carrying allele $i$ that will not mate successfully due to attempting a same-sex mating $n_i^{ss}$ is

$$n_i^{ss} = x_i^{s'}(1 - a_i s)(1 - \sigma). \tag{13}$$

The $n_i^{ss}$ will not be accounted for further in the current reproductive bout as they cannot reproduce successfully.

We assume that each mating produces equal reproductive output and that there is sufficient reproductive excess to keep the population size fixed. As such, the proportion of $A_1$ offspring produced in the population is

$$N_1 = \frac{p_{11} + \frac{1}{2}(p_{12} + p_{21})}{\sum_{i=1}^{2}\sum_{j=1}^{2} p_{ij}}. \tag{14}$$

Likewise, the proportion of $A_2$ offspring is

$$N_2 = \frac{p_{22} + \frac{1}{2}(p_{12} + p_{21})}{\sum_{i=1}^{2}\sum_{j=1}^{2} p_{ij}}. \tag{15}$$

From equations 14 and 15 it can be seen that adding parameters such as $f$, $r$, and $p$ (described above) will have no effect on the model as each pair experiences them equally and they will perfectly cancel. They have thus been ignored. New recruits are then added to the population relative to the proportions $N_1$ and $N_2$. Doing so returns the system to the population size of 1, so the $x_i$ are again frequencies.

The resulting dynamical equations are too complex for meaningful analytical work, however they can readily be analyzed numerically. In particular, evolutionary stable strategies can be found. To do so, we use the initial conditions of $x_1^s = x_1^t = 0.49$ and $x_2^s = x_2^t = 0.01$ and determine whether the rare allele can invade and replace the resident. Starting with steps of 0.1 between $a_1$ and $a_2$ and using successively smaller steps, one can determine the ESS with an arbitrary degree of accuracy. We use $10^{-4}$ in the analysis here. We also test for the possibility of a stable polymorphism by checking for mutual invasibility with starting frequencies of 0.99 and 0.01 for the common and rare allele, respectively. Below we modify the assumptions of the model to check the generality of our conclusions.

#### Additional cost to SSB

As a first test for the robustness of our results, we considered the possibility that same-sex matings carry costs beyond those of opposite-sex matings. To do so, we defined $\delta$ to be the probability that an individual died as a result of a same-sex mating when it would not have died following an opposite-sex mating. This cost can be added to the probability that an individual survives to the next breeding bout $\rho$ (equation (2)). Now, $\rho$ becomes

$$\rho = 1 - (d + ac + f\delta(1-\zeta)). \tag{16}$$

The final term $f\delta(1-\zeta)$ is simply the probability that an individual engages in a same-sex mating ($f(1-\zeta)$) and does not survive to reproduce again as a result. For the rest of this section, we will assume that an individual always finds a potential mate $f = 1$ for simplicity.

The same steps as above can be followed to derive the expected lifetime reproductive success and take the derivative with respect to $a$ to acquire the fitness gradient. As before, we can acquire a meaningful expression for when the fitness gradient is negative at $a = 0$. In this case, sexual discrimination will not be selected for whenever

$$c > -\frac{s(1-\sigma)(d^2 - d(1-2\delta(1-\sigma)) - \delta(1-\delta(1-\sigma)^2))}{\sigma}. \tag{17}$$

By assuming an equal sex ratio ($\sigma = 0.5$), we obtain the simpler condition

$$c > s(d((1-\delta) - d) + (1 - \frac{\delta}{4})\delta). \tag{18}$$

Likewise, the fitness gradient will be positive at $a = 1$ and sexual discrimination will evolve to completion (assuming an equal sex ratio) whenever

$$c < \frac{-(1 + 2ds + s\delta(1-s)) + \sqrt{1 + 4ds(1+s) + 2\delta s(1+s)}}{2s}. \tag{19}$$

The results of this model are compared to the model in the main text in Supplementary Appendix S1.

Finally, a corresponding population genetic model was built. This model is equivalent to the one presented above, however, $\delta$ of the $n_i^{ss}$ are assumed to die between time steps.

#### Multiplicative survival cost

Above, we assumed that the cost to sexual discrimination was paid simultaneously with other forms of mortality. That is, the overall death rate $(1 - d - ac)$ is additive. This could be argued to be likely, however, one could also imagine that other forms of mortality occur sequentially (at a different time point) than mortality due to searching. Then, the overall mortality would be $(1 - d)(1 - ac)$. Now, $d$ is the proportion of the total population that dies due to causes other than searching and $ac$ is the remaining proportion of individuals that die due to searching that survived other sources of mortality (note the reverse order of mortality is mathematically equivalent). Given these assumptions and as explained above, the fitness gradient can be calculated to be

$$\frac{dR_0}{da} = -\frac{(1-d)f(1-r)(1-ps)(a^2c^2(1-d)s(1-\sigma) - ds(1-\sigma) + 2acds(1-\sigma) + c\sigma)}{(d + a(c - cd))^2}. \tag{20}$$

Once more, the sign of the fitness gradient is determined entirely by the third term in equation (20). The sign of the fitness gradient given no attempt to discriminate is again particularly enlightening. In particular, it can be shown from equation (20) that the optimal strategy is to mate entirely indiscriminately whenever

$$c > \frac{ds(1-\sigma)}{\sigma}. \tag{21}$$

Of course, if the sex ratio is equal ($\sigma = 0.5$), this condition simplifies to $c > ds$.

Finally, maximum discrimination will evolve whenever the third term evaluated at $a = 1$ is negative. Assuming an equal sex ratio ($\sigma = 0.5$), maximum attempted discrimination is the best strategy whenever

$$c < \frac{-(1+2ds) + \sqrt{1 + 4ds(1+s)}}{2s(1-d)}. \tag{22}$$

We discuss the biological interpretation and implications of equations (21) and (22) in Supplemental Appendix S2. We also build a population genetic model making the corresponding assumptions. The only difference between the population genetic model described above and the model with multiplicative death, is that a proportion $(1-d)(1-ac)$ die in this version of the model. Again, the results of this model align closely with the optimization approach and are shown in Supplemental Appendix S2.

#### Cost to fecundity

To further test the generality of our hypothesis that indiscriminate mating can be an optimal strategy, we also built a version of the model where the cost of sexual discrimination is carried in fecundity rather than survival (so all individuals die with probability $d$ between each breeding bout). Thus, each mating produces $(1 - ps - ac)$ offspring relative to a mating between an individuals that neither signal nor attempt to discriminate. Note that this gives the additional constraint that $p < 1 - c$, which assumes that $a$ and $s$ are maximized and thus holds $\forall a, s$.

The corresponding fitness gradient from this model is

$$\frac{dR_0}{da} = \frac{(1-d)f(1-r)((1-2ac)s(1-\sigma) - ps^2(1-\sigma) - c\sigma)}{d}. \tag{23}$$

Once more, a simple condition for a negative fitness gradient in the absence of the attempt to discriminate can be derived to determine when the optimal strategy will be to never attempt to discriminate. Now, indiscriminate mating is the optimal strategy whenever the final term is negative, that is

$$c > \frac{s(1-sp)(1-\sigma)}{\sigma}. \tag{24}$$

Or, given an equal sex ratio ($\sigma = 0.5$), $c > s(1 - sp)$.

Similarly, complete discrimination is the best strategy when the fitness gradient is positive at $a = 1$. Assuming an equal sex ratio, this occurs whenever

$$c < \frac{s(1-ps)}{1+2s}. \tag{25}$$

We discuss the biological interpretation and implications of equations (24) and (25) in Supplemental Appendix S3.

Finally, we built a corresponding population genetic model. In this model, all individuals die between breeding bouts with probability $d$. The only required changes are to equations (14) and (15). Matings involving $x_1^s$ (i.e., $p_{11}$ and $p_{12}$) are weighted by $(1 - sp - a_1c)$ and matings involving $x_2^s$ (i.e., $p_{21}$ and $p_{22}$) are weighted by $(1 - sp - a_2c)$. The results from the population genetic model again align closely with the optimization approach and are presented in Supplemental Appendix S3.

#### Semelparous life history

Finally, to determine the importance of life history to our results, we considered a case where the searching sex is semelparous (can only reproduce once). Presumably, more restricted chances to reproduce will disfavor the evolution of indiscriminate mating. This model is a simplification of the first model presented in the sense that the lifetime reproductive succes $R_0$ is now given by equation (3) without the infinite sum.

The resulting fitness gradient is

$$\frac{dR_0}{da} = f(1-ps)((1-2ac-d)s(1-\sigma) - c\sigma). \tag{26}$$

This equation is linear w.r.t. $a$ and can be set to 0 and readily solved for the optimal values of $a$. We see, in particular, that the fitness gradient is negative at $a = 0$ and thus no discrimination should evolve whenever

$$c > \frac{(1-\sigma)(1-d)s}{\sigma}. \tag{27}$$

And when the sex ratio is equal ($\sigma = 0.5$) this becomes $c > (1-d)s$.

Likewise, complete sexual discrimination evolves when the fitness gradient is positive at $a = 1$ which requires

$$c < \frac{(1-d)s}{1+2s} \tag{28}$$

given that the sex ratio is equal. The interpretation and implications of this model is discussed in detail in Supplementary Appendix S4.

As before, we built a corresponding population genetic model. Now, this model is identical to that described under the "Population genetic model" section above except it follows non-overlapping generations. That is, equation (14) and 15 now give the frequencies of $A_1$ and $A_2$ individuals in the next generation.

**Supplementary Appendix S1: Additional cost to SSB**

As mentioned in the main text, same-sex matings have been shown in some cases (and are often believed to be)[3] more costly than matings with the opposite sex. Here, we will determine whether indiscriminate mating can be the optimal strategy in spite of such costs. We define $\delta$ to be the probability that a member of the searching sex dies after a same-sex mating when it would have survived otherwise. After incorporating such a cost (and for simplicity assuming an equal sex ratio), it can be shown (see Supplementary Methods) that indiscriminate mating is the optimal strategy whenever

$$c > s\left[d\big((1-\delta)-d\big)+\left(1-{}^{\delta}/_{4}\right)\delta\right]. \quad \text{(S1)}$$

Unsurprisingly, incorporating additional costs to SSB makes the conditions favoring indiscriminate mating more restrictive. However, a wide range of conditions still exist for which indiscriminate mating is the optimal strategy, even with appreciable costs to sexual discrimination (Fig. S1.1). Most importantly, that small costs to SSB lead to relatively minor changes in optimal evolutionary outcomes (Fig S1.2). In other words, the presence of additional costs to SSB does not immediately cause indiscriminate mating to breakdown. Still, appreciable costs to SSB do lead to higher levels of sexual discrimination evolving in much of parameter space. These conclusions also hold in the population genetic version of the model (Fig. S1.3), although adding an additional cost to SSB influences the results less in this model. This difference occurs for two reasons. First, the additional cost is necessarily multiplicative in the population genetic model due to the timing of when mortality events occur. Second, $\delta$ also skews the sex ratio to be biased towards the targeting sex.

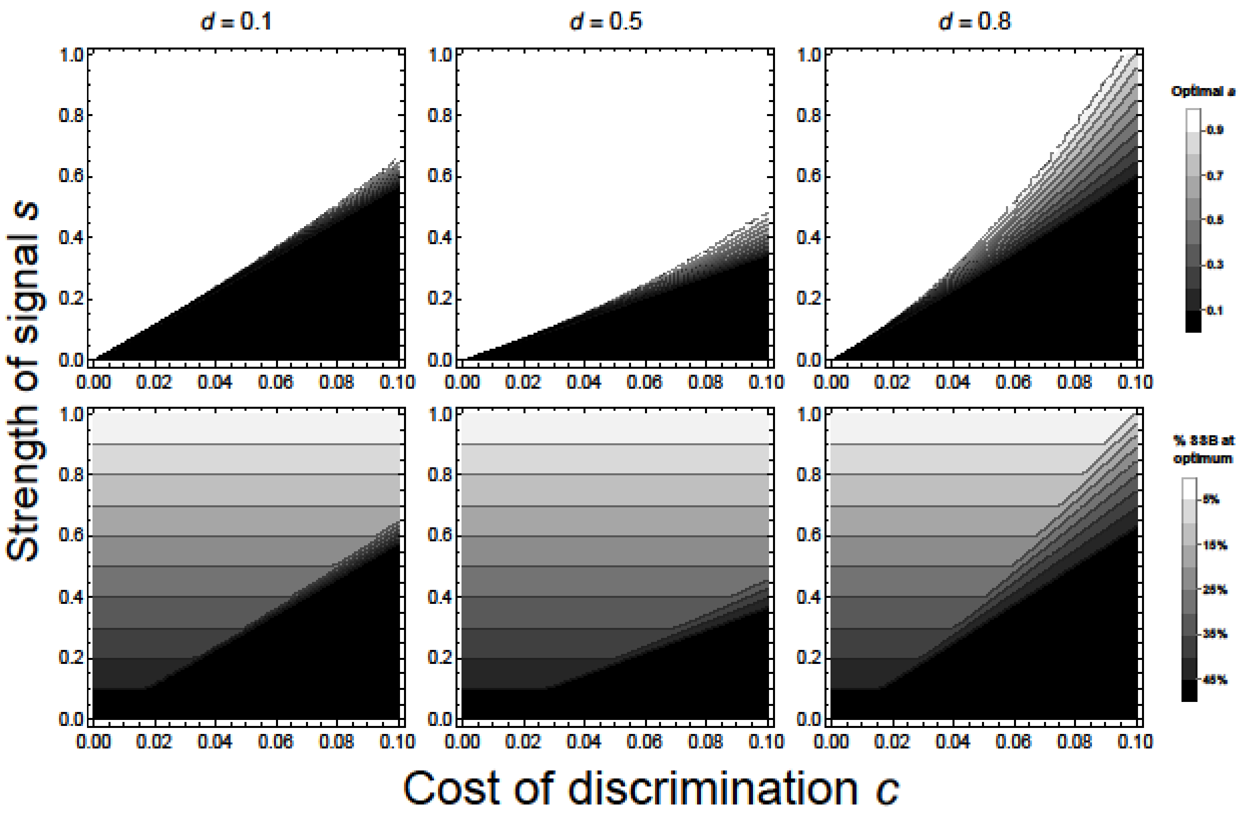


**Fig. S1.1.** Top row: Optimal discrimination strategies (*a*) predicted by the optimization model with an additional cost to SSB with cost to discrimination *c* on the x-axis, strength of sexual signal *s* on the y-axis, and death rate *d* increasing across columns from left to right. Note that the third column has $d = 0.8$ and not $d = 0.9$ as in Fig. 1, as this value is out of range. Bottom row: proportion of matings expected to be SSB at the evolutionary optimum. Darker values indicate more same-sex matings (black: half of matings are with individuals of the same sex if behaving optimally; white: no matings are with individuals of the same sex if behaving optimally). As expected, adding an explicit cost to SSB leads to discriminate mating being the optimal strategy over a wider range of conditions. Other parameters: $\delta = 0.1$, $\sigma = 0.5$, $f = 1$, $r = 0$, $p = 0$.

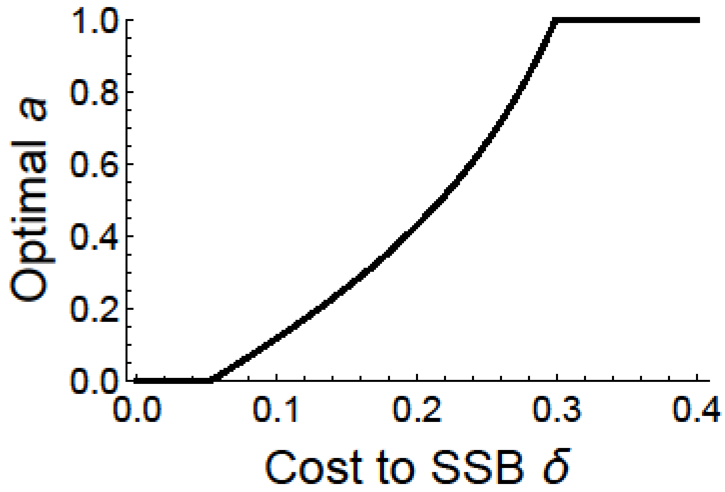


**Fig S1.2.** Optimal sexual discrimination *a* as a function of cost to SSB $\delta$. Here, we see that small changes in the costs to SSB do not qualitatively alter the outcome. Further, specific costs to SSB can drive the optimal strategy from being indiscriminate mating to complete discrimination. Other parameters: $d = 0.7$, $s = 0.4$, $c = 0.09$, $\sigma = 0.5$, $f = 1$, $r = 0$, $p = 0$.

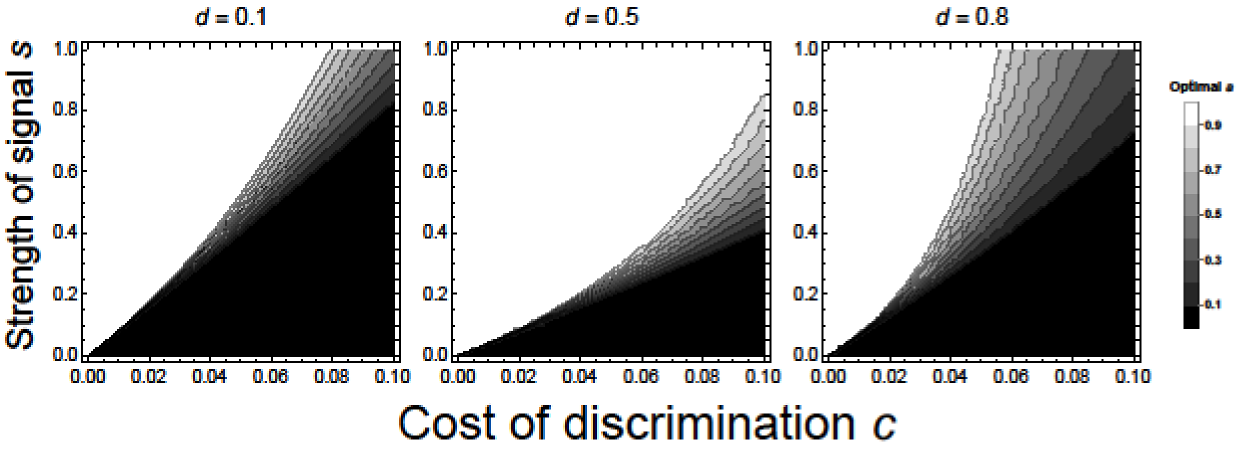


**Fig. S1.3.** ESS level of attempted sexual discrimination from the population genetic model (lighter colors indicate higher attempted discrimination; white: $a = 1$; black: $a = 0$). Again, we see that the ESS level of attempted discrimination increases over some of parameter space (compared to Fig S1), although the effect is minor in the population genetic model. Other parameters: $\delta = 0.1$.

**Supplementary Appendix S2: Multiplicative mortality**

We assume in the main text that death due to sexual discrimination occurs simultaneously with other sources of mortality. Now, we change the form of mortality in our model to allow death due to attempted discrimination and other sources of death to occur sequentially (see Supplementary Methods). This corresponds to background mortality occurring prior to mortality specifically from attempted sexual discrimination (or vice versa). With these assumptions, no sexual discrimination will evolve whenever

$$c > ds(1-\sigma)/\sigma. \quad \text{(S2)}$$

The general conclusions from the original model are maintained, except for the case of high mortality $d$ (Fig S2.1). Now, optimal attempted sexual discrimination increases monotonically with death rate $d$. That is, the largest range of parameter space wherein the best strategy is to mate indiscriminately occurs when the death rate is low. The effect of each parameter in this model is shown in Table S2.1. Once again, our results are supported quite well by the corresponding population genetic model (Fig S2.2).

The key difference between this model and the original model is that now high mortality rates promote sexual discrimination. Unsurprisingly, this can be understood through the multiplicative effect of death. In the original model, the probability of death from sexual discrimination is $ac$. Now, that probability becomes $(1 - d)ac$. Clearly, as $d$ becomes large, the background source of mortality makes it such that mortality from discrimination is essentially negligible. Which model is more realistic depends upon the biology of specific systems.

**Table S2.1**. Summary of parameters, their meaning, and their role in the evolution of sexual discrimination from the model with a multiplicative cost to survival. Note that mortality *d* has the only effect that differs from the main text.

| Parameter | Meaning | Range | Effect |
|---|---|---|---|
| $a$ | Attempt to mate discriminately | $0 \leq a \leq 1$ | N/A |
| $c$ | Cost of mating discriminately | $0 \leq c \leq 1$ | Decreasing $c$ favors sexual discrimination |
| $s$ | Signal (by targeted sex) of their sexual identity | $0 \leq s \leq 1$ | Increasing $s$ favors sexual discrimination |
| $\sigma$ | Proportion of population of the targeted sex | $0 < \sigma < 1$ | Decreasing $\sigma$ favors sexual discrimination |
| $p$ | Cost of presenting sexual identity (by targeted sex) | $0 \leq p < 1$ | Increasing $p$ decreases the selection gradient but has little effect on the ESS |
| $d$ | Baseline mortality probability between each reproductive bout | $0 < d < 1$ | Increasing $d$ favors sexual discrimination |
| $f$ | Probability of finding any individual with which to attempt mating | $0 < f \leq 1$ | Increasing $f$ increases the selection gradient but has little effect on the ESS |
| $r$ | Probability mate of correct sex rejects focal individual | $0 \leq r < 1$ | Increasing $r$ decreases the selection gradient but has little effect on the ESS |

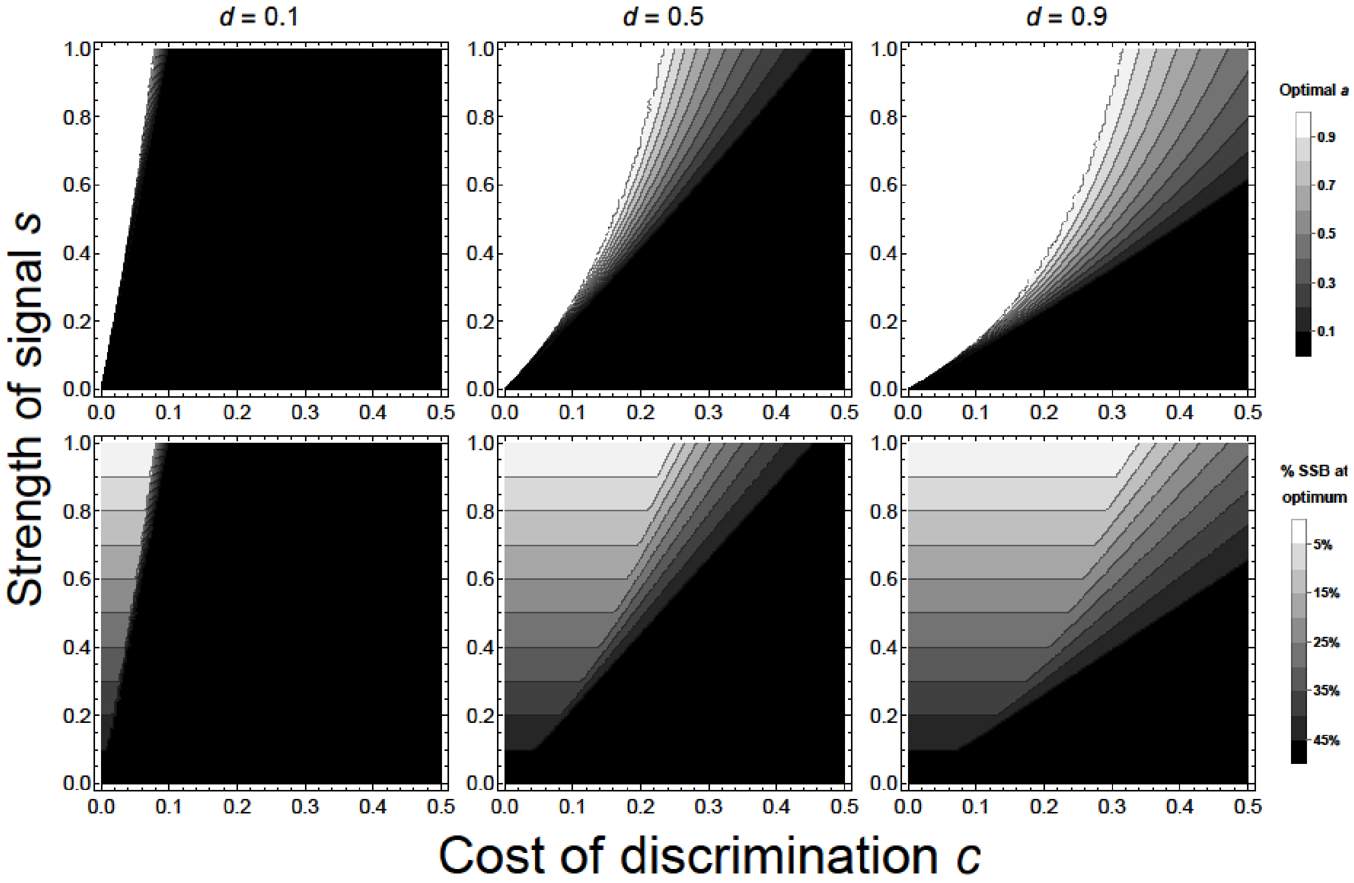


**Fig. S2.1**. Top row: Optimal discrimination strategies (*a*) predicted by the optimization model with multiplicative mortality with cost to discrimination *c* on the x-axis, strength of sexual signal *s* on the y-axis, and death rate *d* increasing across columns from left to right. Note that the x-axis range has expanded from Fig. 1, as separating the sources of mortality changes the acceptable range and meaning of *c*. Higher levels of attempted discrimination correspond to lighter shading (white: *a* = 1; black *a* = 0). Indiscriminate mating is favored by increasing the cost of discrimination *c* or the strength of the sexual signal *s* and decreasing the death rate *d*. Bottom row: proportion of matings expected to be SSB at the evolutionary optimum. Darker values indicate more same-sex matings (black: half of matings are with individuals of the same sex if behaving optimally; white: no matings are with individuals of the same sex if behaving optimally). Other parameters: σ = 0.5, *f* = 1, *r* = 0, *p* = 0.

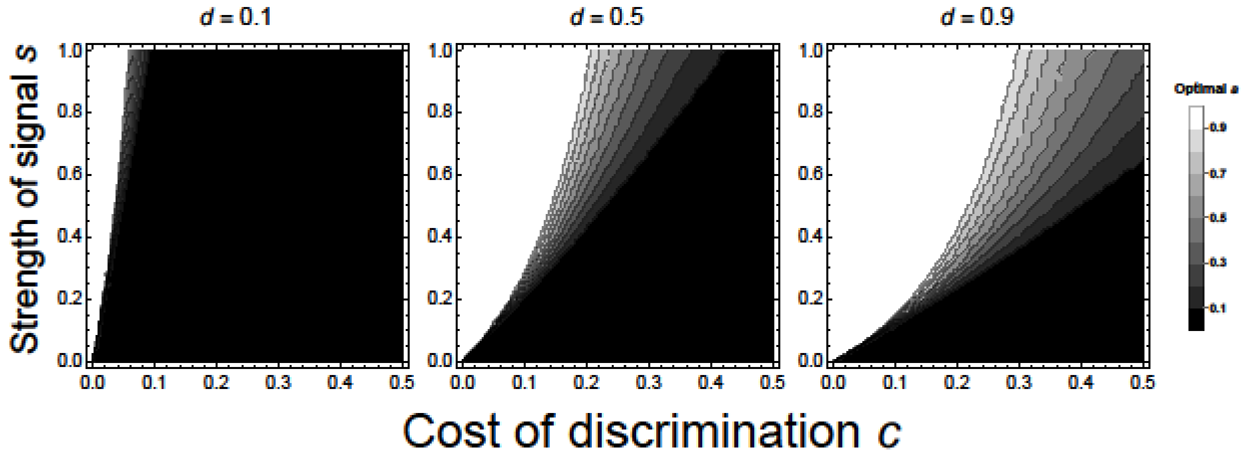


**Fig S2.2**. ESS proportion of attempted sexual discrimination from the population genetic model with multiplicative mortality (lighter colors indicate higher attempted discrimination; white: $a = 1$; black: $a = 0$). Once again, the population genetic model agrees with the method of optimizing expected lifetime reproductive success.

**Supplementary Appendix S3: Fecundity cost to attempted discrimination**

Although search costs are typically considered to be survival costs, one could also build a model wherein the cost to sexual discrimination is to fecundity and not survival. This could correspond to decreasing sperm motility, less energy to put into egg production or egg investment, or less energy to expend on parental care resulting from increasing effort to sexually discriminate. We modify our model to include this assumption in the Supplementary Methods and present the outline of the model and its results here. In this version of the model, we assume that each mating involving an individual that attempted to discriminate produces *c* fewer offspring relative to a mating in which the searching sex did not attempt to discriminate (see Supplementary Methods). Under these assumptions, no sexual discrimination will evolve whenever

$$c > s(1 - ps)(1 - \sigma)/\sigma. \quad \text{(S3)}$$

The conditions for a negative fitness gradient at $a = 0$ in this model are more restrictive than when the discrimination cost is to viability (Fig S3.1). Again, it is predicted that indiscriminate mating is more likely to be the optimal evolutionary strategy when the discrimination cost *c* increases (Fig S3.1). Interestingly, there are some scenarios when the cost to signaling *p* is large in which increasing the strength of the sexual signal *s* actually decreases the optimal level of sexual discrimination *a* (Fig S3.1). This occurs because the benefit of increasing the number of offspring produced in a mating (by attempting to discriminate less; i.e., decreasing *a*) starts to outweigh the benefit of being more likely to mate (by attempting to discriminate more; i.e., increasing *a*) when the cost to signaling *p* is large and *s* is high, resulting in very low overall fecundity. The role of each parameter is described in Table S3.1. Once again, the uninvadable strategies predicted by the optimization approach agree very closely with a corresponding population genetic model (Fig S3.2).

The fact that equations (1) and (S3) differ show the need for empirical work that focuses on determining the costs to sexual discrimination. Although more restrictive, the presence of parameter space in which the preferred strategy is to attempt no sexual discrimination prior to mating demonstrates that SSB as a result of indiscriminate mating can potentially arise in a wide range of ecological and life history scenarios.

**Table S3.1**. Summary of parameters, their meaning, and their role in the evolution of sexual discrimination from the first model presented with discrimination as a cost to survival. Note that only signaling cost *p* and mortality *d* have different effects than in the main text.

| **Parameter** | **Meaning** | **Range** | **Effect** |
|---|---|---|---|
| $a$ | Attempt to mate discriminately | $0 \le a \le 1$ | N/A |
| $c$ | Cost of mating discriminately | $0 \le c \le 1$ | Decreasing $c$ favors sexual discrimination |
| $s$ | Signal (by targeted sex) of their sexual identity | $0 \le s \le 1$ | Increasing $s$ favors sexual discrimination |
| $\sigma$ | Proportion of population of the targeted sex | $0 < \sigma < 1$ | Decreasing $\sigma$ favors sexual discrimination |
| $p$ | Cost of presenting sexual identity (by targeted sex) | $0 \le p < (1-c)$ | Decreasing $p$ favors sexual discrimination |
| $d$ | Baseline mortality probability between each reproductive bout | $0 < d < 1$ | Increasing $d$ decreases the selection gradient but has little effect on the ESS |
| $f$ | Probability of finding any individual with which to attempt mating | $0 < f \le 1$ | Increasing $f$ increases the selection gradient but has little effect on the ESS |
| $r$ | Probability mate of correct sex rejects focal individual | $0 \le r < 1$ | Increasing $r$ decreases the selection gradient but has little effect on the ESS |

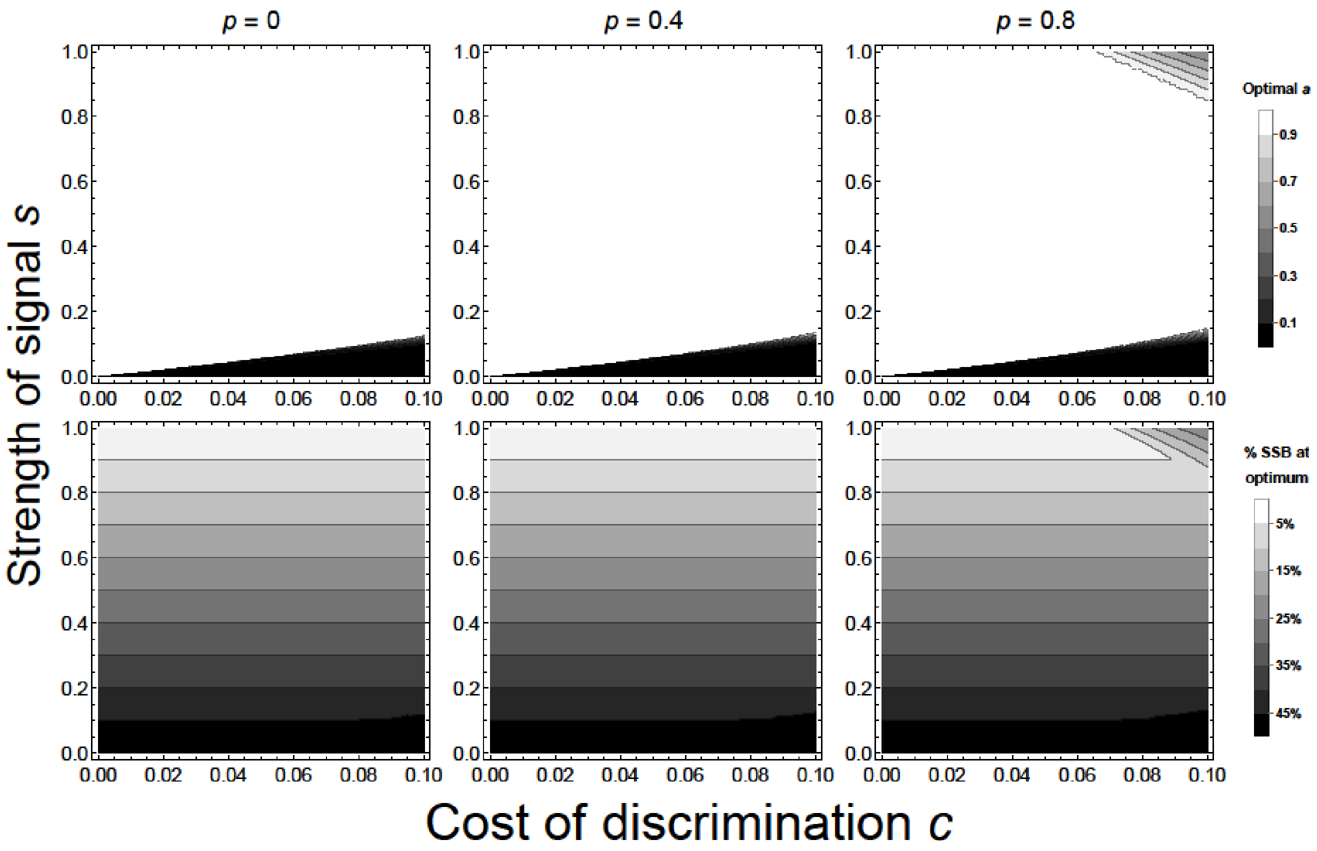


**Figure S3.1.** Top row: Optimal discrimination strategies (*a*) predicted by the optimization model with a discrimination cost to fecundity instead of viability, with cost to sexual signal *p* increasing from left to right. Higher levels of attempted discrimination correspond to lighter shading (white: *a* = 1; black *a* = 0). Indiscriminate mating is favored by increasing the cost of discrimination *c* or the strength of the sexual signal *s* and decreasing the cost to sexual signals *p*. Interestingly when *p* = 0.8, in some cases increasing the sexual signal strength can drive weaker discrimination to evolve. Bottom row: proportion of matings expected to be SSB at the evolutionary optimum. Darker values indicate more same-sex matings (black: half of matings are with individuals of the same sex if behaving optimally; white: no matings are with individuals of the same sex if behaving optimally). Other parameters: $d = 0.5$, $\sigma = 0.5$, $f = 1$, $r = 0$.

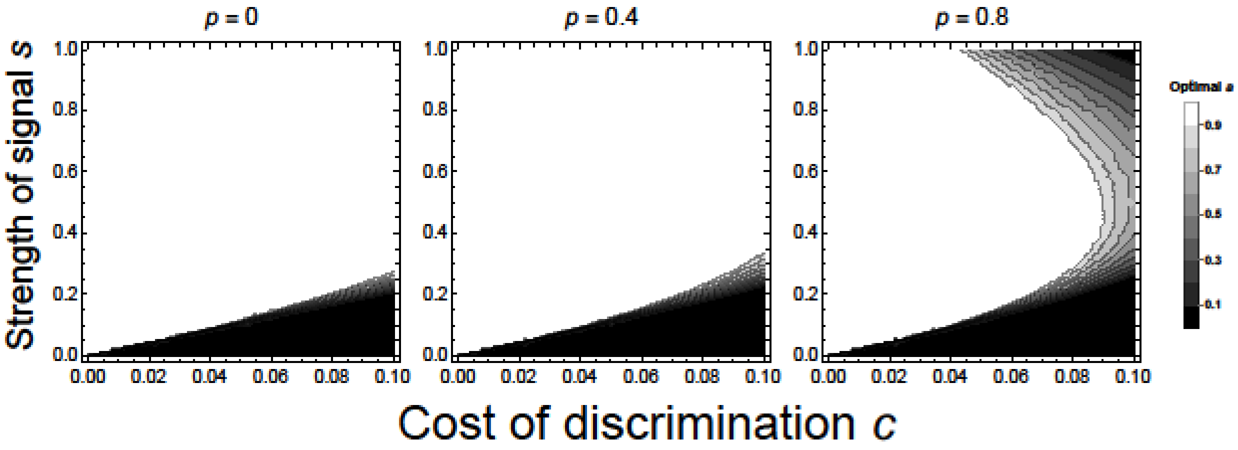


**Figure S3.2.** ESS level of attempted sexual discrimination from the population genetic model with a discrimination cost to fecundity (lighter colors indicate higher attempted discrimination; white: $a$ = 1; black: $a$ = 0). Once again, the population genetic model agrees closely with the optimization model (Fig. S3.1).

#### Supplementary Appendix S4: Semelparous life history

Finally, we considered the case wherein individuals only survived for one reproductive bout (i.e., the searching sex was semelparous). We use the same assumptions from the model presented in the main text with an additive mortality cost to sexual discrimination with the exception that the death rate *d* is now taken to be the proportion of individuals that die prior to attempting reproduction. As individuals have only a single opportunity to reproduce, their fitness includes the term $(1 - d)$ as opposed to the infinite sum from before. Under these assumptions, no sexual discrimination will evolve whenever

$$c > (1 - \sigma)(1 - d)s/\sigma. \quad \text{(S4)}$$

The major difference between this model and the original model with the iteroparous life history is that mortality *d* now has a purely directional effect on the evolution of sexual discrimination. Namely, increasing mortality *d* favors the evolution of indiscriminate mating (Fig. S4.1). This shows the unsurprising effect that low mortality rates can only favor indiscriminate mating when they result in many opportunities to mate (as explained in the main text). The roles of all other parameters are consistent with the iteroparous model and presented in Table S4.1.

Once again, the assumptions of this optimization model were converted into a population genetic model. The results of these two models align closely as seen in Fig. S4.2.

**Table S4.1**. Summary of parameters, their meaning, and their role in the evolution of sexual discrimination from the first model presented with discrimination as a cost to survival. Again, the only effect that changes from the model in the main text is to mortality *d*.

| **Parameter** | **Meaning** | **Range** | **Effect** |
|---|---|---|---|
| $a$ | Attempt to mate discriminately | $0 \leq a \leq 1$ | N/A |
| $c$ | Cost of mating discriminately | $0 \leq c \leq 1$-d | Decreasing $c$ favors sexual discrimination |
| $s$ | Signal (by targeted sex) of their sexual identity | $0 \leq s \leq 1$ | Increasing $s$ favors sexual discrimination |
| $\sigma$ | Proportion of population of the targeted sex | $0 < \sigma < 1$ | Decreasing $\sigma$ favors sexual discrimination |
| $p$ | Cost of presenting sexual identity (by targeted sex) | $0 \leq p < 1$ | Increasing $p$ decreases the selection gradient but has little effect on the ESS |
| $d$ | Baseline mortality probability prior to the reproductive bout | $0 < d < 1$ | Decreasing $d$ favors sexual discrimination |
| $f$ | Probability of finding any individual with which to attempt mating | $0 < f \leq 1$ | Increasing $f$ increases the selection gradient but has little effect on the ESS |
| $r$ | Probability mate of correct sex rejects focal individual | $0 \leq r < 1$ | Increasing $r$ decreases the selection gradient but has little effect on the ESS |

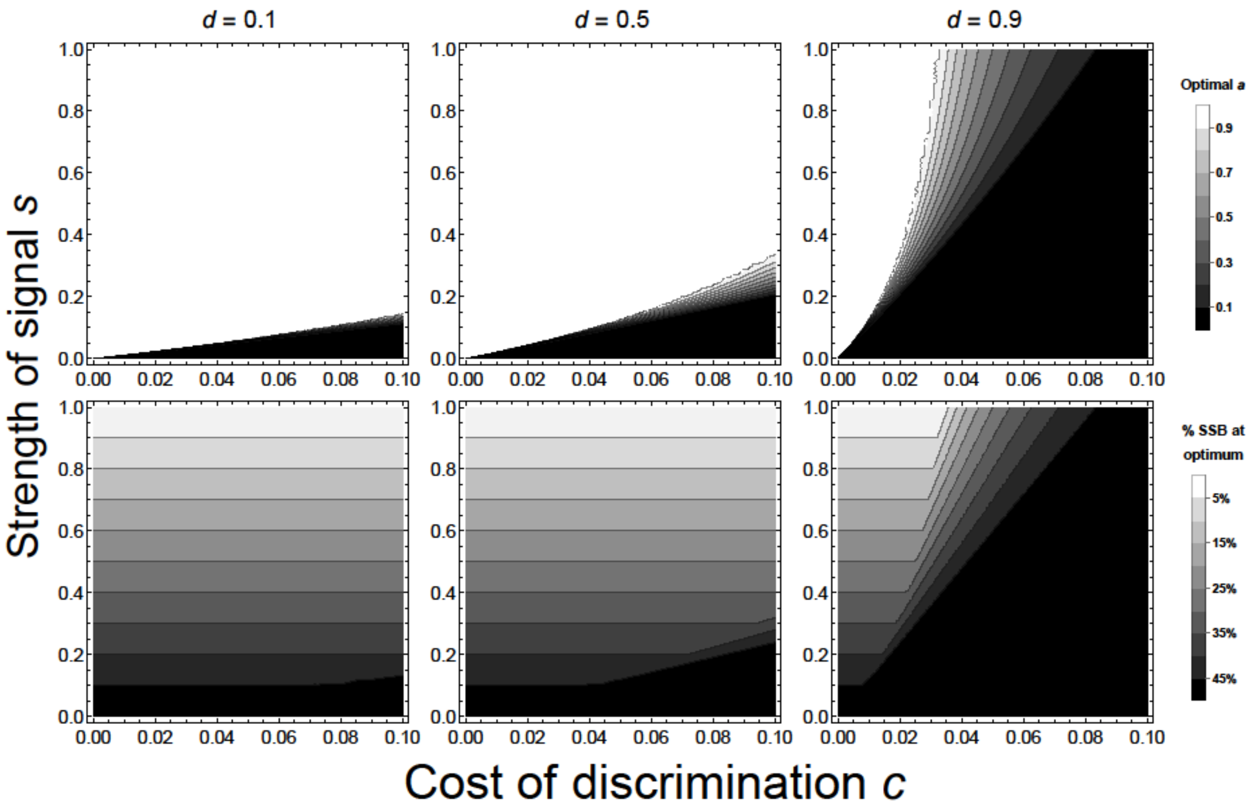


**Figure S4.1.** Top row: Optimal discrimination strategies (*a*) predicted by the optimization model with semelparous life history, with pre-reproductive mortality probability *d* increasing from left to right. Higher levels of attempted discrimination correspond to lighter shading (white: $a = 1$; black $a = 0$). Indiscriminate mating is favored by increasing the cost of discrimination *c* or the strength of the sexual signal *s* and increasing mortality *d*. Bottom row: proportion of matings expected to be SSB at the evolutionary optimum. Darker values indicate more same-sex matings (black: half of matings are with individuals of the same sex if behaving optimally; white: no matings are with individuals of the same sex if behaving optimally). Other parameters: $\sigma = 0.5$, $f = 1$, $r = 0$, $p = 0$.

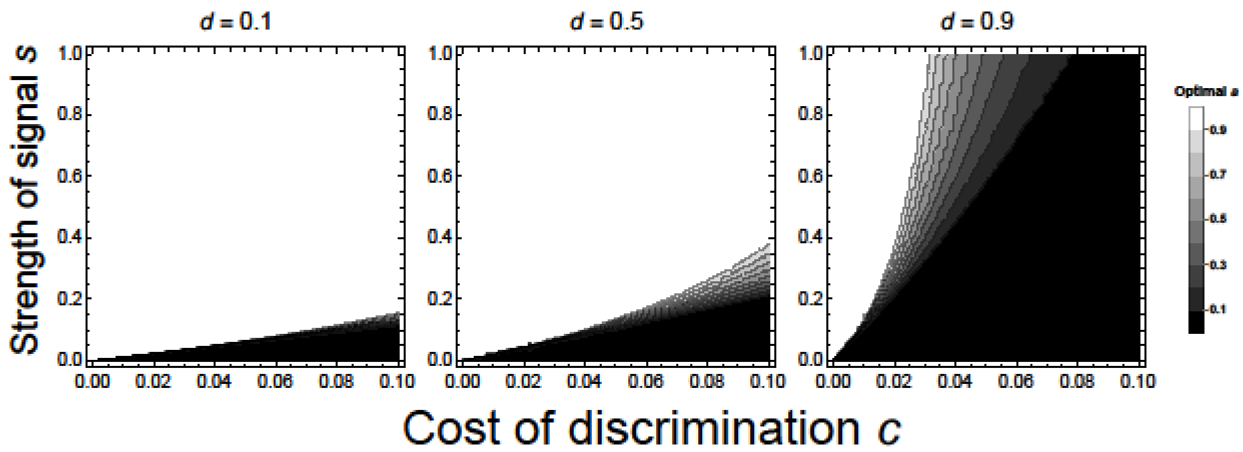


**Figure S4.2.** ESS level of attempted sexual discrimination from the population genetic model with a discrimination cost to fecundity (lighter colors indicate higher attempted discrimination; white: $a$ = 1; black: $a$ = 0). The population genetic model agrees closely with the optimization model (Fig. S4.1).
